# Assessing the role of pleiotropy in the evolution of animal color and behavior: a meta-analysis of experimental studies

**DOI:** 10.1101/2024.01.12.575404

**Authors:** Sarah N. Ruckman, Eve A. Humphrey, Lily Muzzey, Ioanna Prantalou, Madison Pleasants, Kimberly A. Hughes

**Affiliations:** Department of Biological Science, Florida State University, Tallahassee, FL, USA; Biology Department, Lincoln University, Lincoln University, PA, USA

**Keywords:** aggressive behavior, badge of status, coloration, condition dependence, pleiotropy

## Abstract

Color varies in pattern and degree across the tree of life. In animals, genetic variation in color is hypothesized to have pleiotropic effects on a variety of behaviors, due to shared dependence on underlying biochemical pathways. Such pleiotropy can constrain the independent evolution of color and behavior. Although associations between color and behavior have been reported, this relationship has not yet been addressed across a broad taxonomic scale with a formal meta-analysis. We used a phylogenetic meta-analytic approach to examine the relationship between individual variation in aggressive behavior and variation in multiple colors. Seventy studies met our inclusion criteria (vertebrates = 66; invertebrates = 4). After accounting for phylogeny and correcting for publication bias, there was a positive association between measures of aggression and degree or area of coloration (mean = 0.274, 95% CI = (0.041, 0.481)). However, this positive association was not influenced by type of color or by several other variables that we tested. Because the data supports a positive association between aggression and degree or area of coloration, irrespective of whether color is melanin-based, carotenoid-based, or structural, we conclude that this pattern does not strongly support the melanin-pleiotropy hypothesis. The relationship was also unaffected by moderators accounting for individual condition, social rank, or age; thus, the results do not strongly support hypotheses that condition-dependence accounts for relationships between color and aggressive behavior. We propose that the moderate positive correlation between aggression and coloration across *Animalia* that we observed is underlain by genetic covariation between behavior and color traits that serve as badges of status.

## Introduction

A key goal of modern evolutionary biology is predicting if and how populations will evolve adaptively in response to environmental change. Predicting adaptive evolution requires knowing if there are constraints on how populations can respond to selection. The definition of adaptive constraint has been debated, but the unifying theme of these definitions is that populations are not always able to respond to selection in predicable ways (Blows and Walsh 2009; Walsh and Blows 2009). Pleiotropy, and the genetic covariance between traits that can result, is one potential cause of adaptive constraint. However, estimating genetic covariance in natural populations requires very large sample sizes and reliable information on relatedness which are currently available in few species. Trait covariance can also be environment-dependent, meaning many such estimates might be necessary to accurately predict responses to selection in variable environments.

Another approach is to use functional genetic information to predict pleiotropic effects and their population-level consequences. A well-known example is the hypothesis that, in animals, the biochemical pathways that regulate body coloration and behavior can overlap, leading to genetic covariance between color and behavior. Specifically, in vertebrates, melanic coloration arises from the binding of melanocortin agonists to the G-protein-coupled receptor MCR-1. Melanocortins (peptide hormones derived from the prohormone Proopiomelanocortin) also bind to other receptors that regulate diverse functions, including aggressive behavior (Ducrest et al. 2008). Therefore, one prediction of this hypothesis is that heritable variation in abundance or activity of melanocortins generates genetic correlation between body color and aggression; animals with higher levels of eumelanic (dark brown, black) color are expected to exhibit higher levels of aggression (Ducrest et al. 2008; Roulin and Ducrest 2011). We note that other kinds of changes in the melanocortin system, such as mutation in melanocortin receptors that are expressed both in the skin and the brain, could also produce these correlations (e.g., Reissmann and Ludwig 2013). In support of this hypothesis, studies have reported correlations between melanin-based coloration and behavior, including aggressive behavior, in vertebrates (e.g., Moore and Martin 2016; Dijkstra et al. 2017; Seddon and Hews 2017; Beck et al. 2018). This hypothesis has recently been extended to include insects and other invertebrates (San-Jose and Roulin 2018), where melanin is synthesized from dopamine. This invertebrate biosynthetic pathway provides a plausible link between body coloration and behavior (Wittkopp and Beldade 2009; Massey and Wittkopp 2016), and predicts correlations between traits that are similar to those seen in vertebrates have been reported (San-Jose and Roulin 2018).

Other functional links between color and behavior have been hypothesized to arise from carotenoid condition dependence, whereby animals in good condition can devote more energy to high levels of aggression, and to carotenoid pigment synthesis, which is dependent on obtaining carotenoids in the diet (Blount and McGraw 2008; Backström et al. 2015). Under this scenario, genetic variants affecting foraging ability would simultaneously affect carotenoid color and aggressiveness, leading to genetic correlations between these traits. However, carotenoid condition-dependence could also arise from purely environmental covariance between aggression and color if, for example, individual variation in foraging success arises from non-genetic variation (e.g., variation in habitat quality). Purely environmental covariance between color and aggression would not impose the same kind of adaptive constraint as would a genetic correlation.

A third hypothesis is that a broad range of colors can be indicators or “badges” of social status. Badges of status are traits (e.g., color patches) that influence the outcome of aggressive encounters (Rohwer 1975; Dawkins and Krebs 1978). Badges of status are not limited to specific colors; badges can even lack color, as seen in the white forehead patch of collared flycatchers *Ficedula albicollis* (Pärt and Qvarnström 1997). A meta-analysis of associations between dominance and plumage characteristics (color, UV presence, and color patch size) in birds reported a positive correlation between dominance and measures of coloration, irrespective of specific color; the authors interpreted this result as supporting the badge of status hypothesis (Santos et al. 2011; but see Sánchez-Tójar et al. 2018). A comparative analysis of competing bird species, found that dominant species have on average more black than subordinate species; carotenoid and other colors were sometimes, but not always associated with dominance (Kenyon and Martin 2023). The authors interpreted this result as also supporting the badge of status hypothesis (Kenyon and Martin 2023). Such badges can be honest signals (reliable predictors of aggressiveness), although the mechanism maintaining honesty in the signal has been debated and could be different for each color or type of badge (Johnstone and Norris 1993; Tibbetts and Dale 2004). In ruffs, *Philomachus pugnax,* a supergene relates color and aggression, where the genes associated with color and behavior are located near each other and in linkage disequilibrium (Küpper et al. 2016; Lamichhaney et al. 2016). In contrast, some badges of status might not vary genetically, but might instead vary due to non-genetic sources. For example, some badges are plastic in expression and can vary as a result of dominance interactions (e.g., Dey et al. 2014). As above, implications for adaptive constraint depend upon the underlying cause of trait covariance.

Other hypotheses can be consistent with a positive association between aggression and color, irrespective of the type of color. For example, although melanin is endogenously produced, production and/or maintenance of melanic body color might be condition-dependent, although the empirical evidence for this is mixed (Roulin 2016). Similar arguments can be made for structural colors. If structural color depends on the condition of feathers or scales, then costs could be incurred in maintaining or cleaning those structures. Structural color variation has been associated with condition (e.g., McGraw et al. 2002) and mate quality (e.g., Siefferman and Hill 2003) in birds. If aggressive behavior is also condition-dependent, then a general condition dependence might be responsible for positive associations between color and aggressive behavior, irrespective of type of color.

If color and behavior are genetically linked via pleiotropy under any of the above scenarios, independent expression and evolution of these traits can be constrained. This constraint has mainly been discussed in relation to the melanocortin hypothesis, perhaps because melanocortin synthesis is less likely to be condition dependent than other pigment-based colors (Ducrest et al. 2008; Roulin and Ducrest 2011; San-Jose and Roulin 2018). However, counterexamples to the expected association between darker color and higher aggression have been reported. Boerner and Krüger (2009) found that, in the common buzzard (*Buteo buteo*), light colored males are more aggressive than darker colored birds. In pied flycatchers (*Ficedula hypoleuca*), no significant relationship between male color and aggressive behavior was found (Huhta and Alatalo 1993). Examples such as these suggest that species (or populations within species) vary in the relationship between color and aggression. Counterexamples also raise the possibility that the preponderance of studies reporting significant correlations (Ducrest et al. 2008; Roulin and Ducrest 2011; San-Jose and Roulin 2018) reflects taxonomic or publication bias, or that correlations arise from specific features of studies like the age, sex, or condition of the focal animals. For example, a meta-analysis reported positive correlation between dominance and plumage traits in birds; this correlation was unaffected by the type of plumage trait, but was influenced by the assessment method (whether aggression was assessed by quantifying specific aggressive acts, or by an indirect method such as distance between individuals; Santos et al. 2011). We know of no meta-analyses of associations between coloration and behavior that extend across broader taxa, or that assess other factors such as whether color is fixed or plastic during adulthood.

Here, we describe a meta-analysis of the (within-species) relationship between body color variation and aggressive behavior across a broad taxonomic scale. The meta-analysis included vertebrate and invertebrate taxa and controlled for effects of phylogeny on statistical inference. We investigated the possibility that publication bias has influenced the patterns reflected in the published literature. In addition, we evaluated whether the relationship between color and aggression is moderated by the type of color class (e.g. eumelanic, carotenoid, or structural), whether coloration is fixed or varies plastically during adulthood, life stage and sex of the focal animals, type of population studied (wild, domestic, lab-reared, or wild caught and then lab tested), type of aggressive act measured (direct or indirectly measured), geographic origin of the species or source population of the focal animals, and whether or not social rank, age, and the condition of the animal were controlled or measured. We were especially interested in whether associations were moderated by color type because the melanocortin pleiotropy hypothesis predicts specifically that variation in eumelanin-based colors should be associated with aggression. By contrast, condition-dependence has most often been discussed in relation to carotenoid pigmentation (and predicts a positive association between carotenoid pigmentation and aggression). Consequently, positive associations between color and aggression that are restricted to melanic or carotenoid colors would provide direct support for those hypotheses. Both the badge-of-status hypothesis and general condition-dependence predict that associations should be found between aggression and color, but that these associations should not be limited to melanin- or carotenoid-based colors. The general condition dependence mechanism depends upon both color and aggression being condition dependent, leading us to predict that controlling for condition should moderate the relationship between color and behavior. By contrast, the badge-of-status hypothesis does not, *a priori*, make that same prediction.

## Methods

### Data Collection

We defined aggression as any variable that measured antagonistic behaviors (e.g., biting, or chasing) toward a conspecific (of same sex and age) or mirror image. We narrowed our search to conspecifics because sexes may differ in both behavior and coloration as well as how they respond to differing individuals (e.g., Horth 2003). The colors included those regulated by melanin-, carotenoid-, and pterin-based pigments. We also included colors produced by structural variation in skin, scales, feathers, and cuticles (e.g., blue, purple, and white colors produced by iridophores in some fish; Schartl et al. 2016). Each color classification per species was listed and justified with citations in Supplemental Table S1.

We searched Web of Science using the search terms “color” AND “aggression” OR “aggressive” (Figure 1). This yielded 751 papers as of September 2020. We then filtered the list by removing papers with human subjects (n = 197). We removed human subjects because the variation in human pigmentation is not due to the same mechanisms that have been proposed to lead to pleiotropic effects on behavior (Deng and Xu 2017; San-Jose and Roulin 2018). We also removed any papers that did not measure both individual color and aggression (n = 324), which is necessary to determine an association. This yielded 230 papers. Finally, we removed papers if they did not include any of the following: natural color variation within the same species, individual measures of both color and aggression against a con-specific (same age, sex, and color), report relevant statistics for effect size calculations, and analyze the data in comparison between color morphs. This yielded 70 papers (159 effect size estimates) that were included in the phylogenetic meta-analysis.

**Figure 1:**
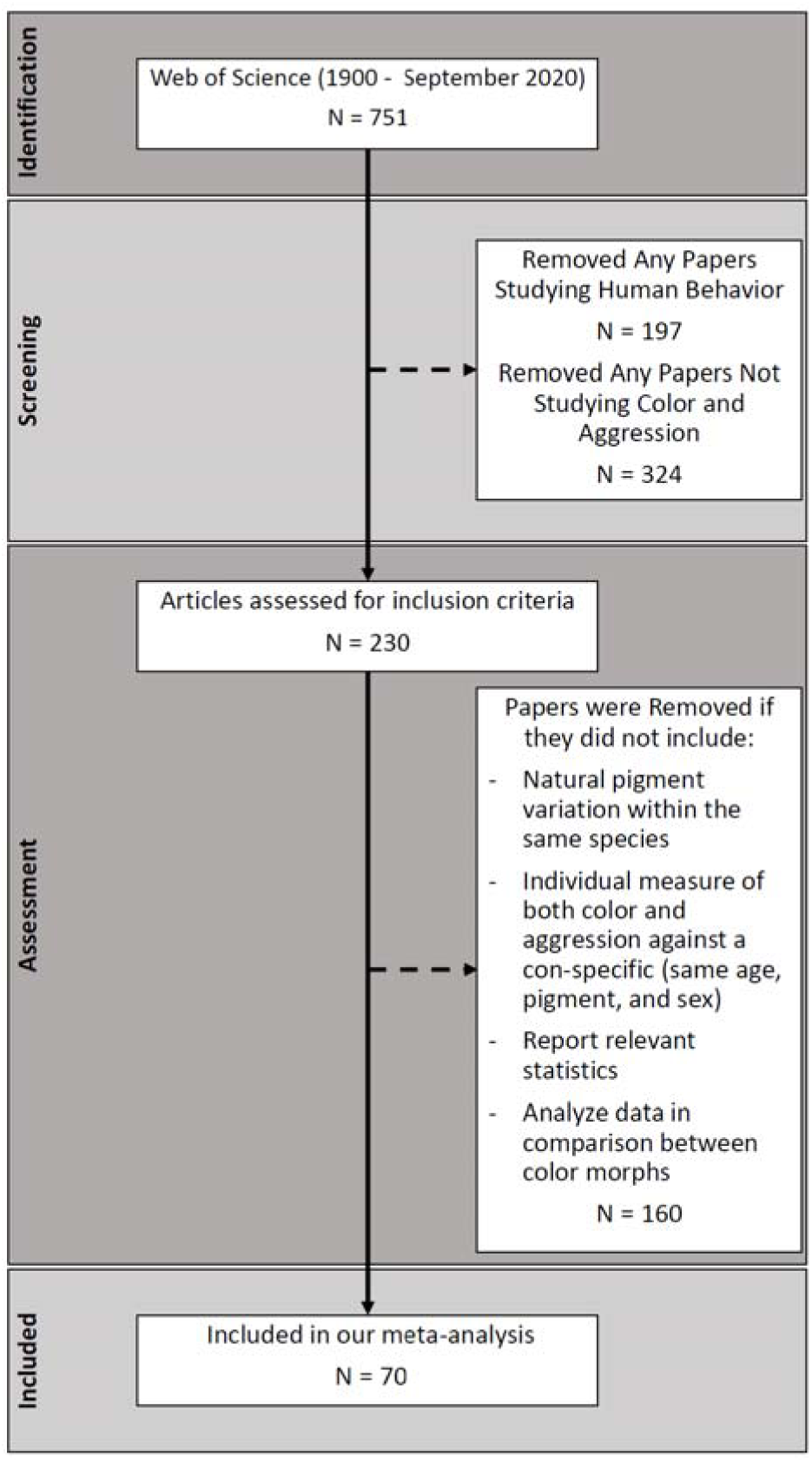
PRISMA Diagram for Papers Examined. Solid arrows indicate papers that moved on to the next step and dashed arrows indicate papers that were removed from the analysis. First, we identified papers using Web of Science and found 751 papers. Next, we screened these papers for human studies and those that clearly did not examine color and aggression. This removed 521 papers. Finally, we assessed the 230 papers based on our inclusion criteria and removed any studies that did not meet all of the inclusion criteria. We found that 70 papers met our inclusion criteria and were analyzed in our meta-analysis.

From each paper, we collected the species name, color (how color was categorized or quantified in the original study, (e.g., white vs. tan, area of eumelanic pigmentation, etc.), location of the color (e.g., total body color or eye color), type of color class (e.g. eumelanin or pheomelanin, carotenoid, structural, pteridine, or unknown), and the measure of aggression used (e.g., direct aggression such as bites, or indirect such as proximity). We also collected the life stage (adult vs juvenile) and sex of the focal individuals, location of the study (wild, lab-reared, domesticated, wild individuals measured in the lab), vertebrate or invertebrate taxa, plasticity of pigment expression (plastic or fixed), seasonality of pigment trait (year-round, breeding, or non-breeding), geographic location of focal or source population, if the condition of the animal was controlled and how (e.g., weight, length, or weight and length), if social rank was considered in the study design and how (e.g., uncontrolled, dummy used, isolated, or recording used), if the age of the animal was controlled and how (e.g., same age or covariate in analysis) and whether the study was an observational or experimental study. Finally, we recorded the measure of association between color and aggression (means/standard deviations of discrete groups, t-, F-, or *χ*^2^-test statistics with the associated p-values and degrees of freedom, or a correlation coefficient), and the sample sizes for each measure of association.

### Effect Size Calculations

For a standardized effect size, when possible, we used reported Pearson’s correlation coefficients between color and aggression, which we then converted to Fisher Z statistics. The Fisher Z transformation is recommended to normalize the sampling distribution of correlation coefficient estimates when sample sizes are small and produces less biased results (Silver and Dunlap 1987; Berry and Mielke 2000). The Fisher Z transformation also widens the distribution around 0, which is useful because Markov Chain Monte Carlo’s (MCMC) have difficulty producing accurate estimates when the true value of the mean is very close to zero (Lipsey and Wilson 2001; Hadfield 2010).

When studies did not report correlation coefficients, we used reported F or *χ*^2^ statistics and converted these into correlation coefficient estimates using methods described in Nakagawa et al. (2007) and Lipsey and Wilson (2001). When studies analyzed categorical data using a t test or reported only means and standard deviations for discrete groups, we could not calculate the product-moment correlation coefficient (Pearson’s r) and instead calculated the biserial correlation coefficient (Jacobs and Viechtbauer 2017). The biserial-correlation coefficient is comparable to the product-moment correlation and can therefore be used in the same meta-analysis (Jacobs and Viechtbauer, 2017). We then converted both Pearson’s and biserial correlation estimates to Fisher Z using the R package *DescTools* and the command FisherZ (Signorell 2021). Biserial-correlation coefficients are sometimes calculated to be greater than 1 or less than -1 (Pustejovsky 2014); values outside this range are undefined under the Fisher Z transformation (Fisher 1915; Silver and Dunlap 1987). This was the case with five positive values in our dataset; these five positive values were reasonably close to 1, total range = -0.77 to 1.33. We therefore converted the positive values to 0.99, as appropriate, as recommended by Pustejovsky (2014).

### Phylogenetic Meta-Analytic Model

To account for the non-independence of the data due to evolutionary history and the relationship between species, we constructed a phylogenetic tree of all the species (70 in total, 66 vertebrates, and 4 invertebrates across 9 classes; Supplemental Figure S1). We used an ultrametric tree that was fully resolved to the species level, which we obtained from TimeTree.org (Hedges et al. 2006; Kumar et al. 2017). We then rooted the tree with the anemone *Phymactis clematis* as the outgroup because it falls outside of Bilateria, to which all species in our dataset belong. We used the R packages *ape, phytools,* and *TreeTools* to obtain the relatedness matrix, root the tree, change the edge lengths from 0 to 0.00001, and plot the tree (Paradis et al. 2004; Revell 2012; Smith and Wickham 2019).

We used the R package *MCMCglmm* to perform the meta-analysis (Hadfield 2010). We chose an expanded prior with a Cauchy distribution that mirrored the Fisher Z distribution (Adams 2008). We ran each model for 2,000,000 interactions and removed 1,000,000 steps as the burnin with a thin of 1,000. Once we determined the models with the lowest DIC values, we reran the analysis using 5,000,000 iterations with burnin of 2,500,000 and a thin value of 1,000; these parameters always produced convergence and final values that were in the Fisher Z distribution bounded by [-2.64, 2.64]. All credible intervals reported are based on the last 2,500,000 iterations of the *MCMCglmm*. Prior to all analysis, we removed one data point that was the sole value associated with the color class pteridine (Robertson and Rosenblum 2010).

### Random and Mixed Models

We first evaluated a model containing only random effects of species, study, the weights associated with each study (calculated as the inverse of the standard error for the Fisher Z for each study), and the phylogenetic tree. Weights were added as a variance-covariance matrix using the “us” option; phylogenetic information was incorporated as a relatedness matrix with the “pedigree” option, which assumes a Brownian-motion model of evolution (Hadfield 2010; Nakagawa and Santos 2012). We removed the random effects using backward elimination to confirm the importance of each random effect. We found that the model with the lowest DIC value was that which included all random effects (see Results). Retaining all random effects, we then assessed whether any moderators improved model fit compared to the random-effects-only model (hereafter, “random effects model”). We tested each of the following moderators one at a time, and in all two-way combinations: type of color classification, plasticity, sex, life stage, vertebrate or invertebrate, location of the study, seasonality of the color, geographic location of the study population, observational vs. experimental studies, if condition of the animal was controlled and how, if social rank was controlled and how, if age of focal animals was controlled and how, and type of aggressive acts (direct or indirect). We also tested for two-way interactions between pairs of moderators that we deemed biologically likely: color class by plasticity, color class by sex, and sex by plasticity. We compared these models using DIC values. For the best-fitting models, we computed medians and 95% confidence intervals of the effect size for each moderator using the R package *emmeans* (Lenth et al. 2021).

Some of the moderators listed above were not available in all the studies in our data set: condition, social rank, and age of focal animals. We accounted for this in two ways. First, as described above, we used a moderator that indicated whether the feature was controlled for or used as a covariate in the study. For example, social rank could be uncontrolled, controlled by using unfamiliar animals, or controlled by using a mirror test or video. For each of these three moderators, we also asked if including only those studies that controlled for the moderator produced substantially different results. Because we were particularly interested in any difference in effect size between different color types, we also included color class as an additional moderator in these “subset” models.

Finally, we investigated whether effect sizes that were reported for “unknown” color classes were potentially modulated by the melanocortin pathway. Recent studies suggest that, at least in fish, the melanocortin system can regulate non-melanin-based colors (reviewed in Cal et al. 2017). To determine if expanding the color classes regulated by the melanocortin system affected the results, we recoded all unknown color classes as eumelanin and re-ran the mixed effects model with color class as a moderator.

### Heterogeneity

Ecology and evolutionary biology meta-analyses are likely to exhibit high levels of heterogeneity due to differences in taxa and experimental methods across studies (Gurevitch and Hedges 1999; Senior et al. 2016). We used a method developed by Nakagawa and Santos (2012) to quantify the proportion of the total variance due to phylogeny (termed “phylogenetic signal” or H^2^) their Equation 26, and the heterogeneity due to study 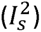 and species 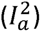 using Equations 24 and 25 of that paper, respectively. To make the values more easily interpretable, we reported the heterogeneity values as percentages, rather than proportions. Percentages close to 0 indicate low heterogeneity while percentages close to 1 are considered high heterogeneity. The phylogenetic signal was left as a proportion.

### Publication Bias

Publication bias due to selective reporting of results can influence both the estimated magnitude and reliability of the overall effect size estimate (Rosenthal 1979; Gurevitch and Hedges 1999). We investigated the possibility of publication bias by visualizing study asymmetry using contour enhanced funnel plots and by testing for asymmetry using a modified Egger’s test as described in Nakagawa and Santos (2012). This test regresses the meta-analytic residuals on the precision of each effect size estimate (inverse of standard error). These residuals, unlike the weighted effect sizes themselves, are independent, and thus satisfy the assumptions of the test. We corrected for the effect of publication bias using the trim-and-fill method implemented in the R package *meta* (Duval and Tweedie 2000; Schwarzer 2007). This method restores funnel plot symmetry by iteratively removing studies with large positive residuals and imputing missing effect size estimates. Because we used meta-analytic residuals for this analysis, the resulting estimates of the mean effect size estimates are the adjustment required to restore the funnel plot symmetry (Duval and Tweedie 2000). All analyses were conducted in R version 4.3.1 (R Core Team 2023). Data and scripts for all analyses are available in a GitHub repository (https://github.com/sruckman/meta-analysis).

## Results

### Full data set analysis

We calculated 159 effect sizes from 70 studies that met our inclusion criteria. The random effects model with the lowest DIC value included all random effects (phylogenetic tree, species, study, and weights; Supplemental Table S2). This model had a mean posterior effect size of 0.251 (95% credible interval = (0.018, 0.458); Pearson’s correlation mean = 0.246 and 95% credible interval = (0.018, 0.428)), indicating support for a positive association between the intensity of color and aggression (Figure 2).

**Figure 2:**
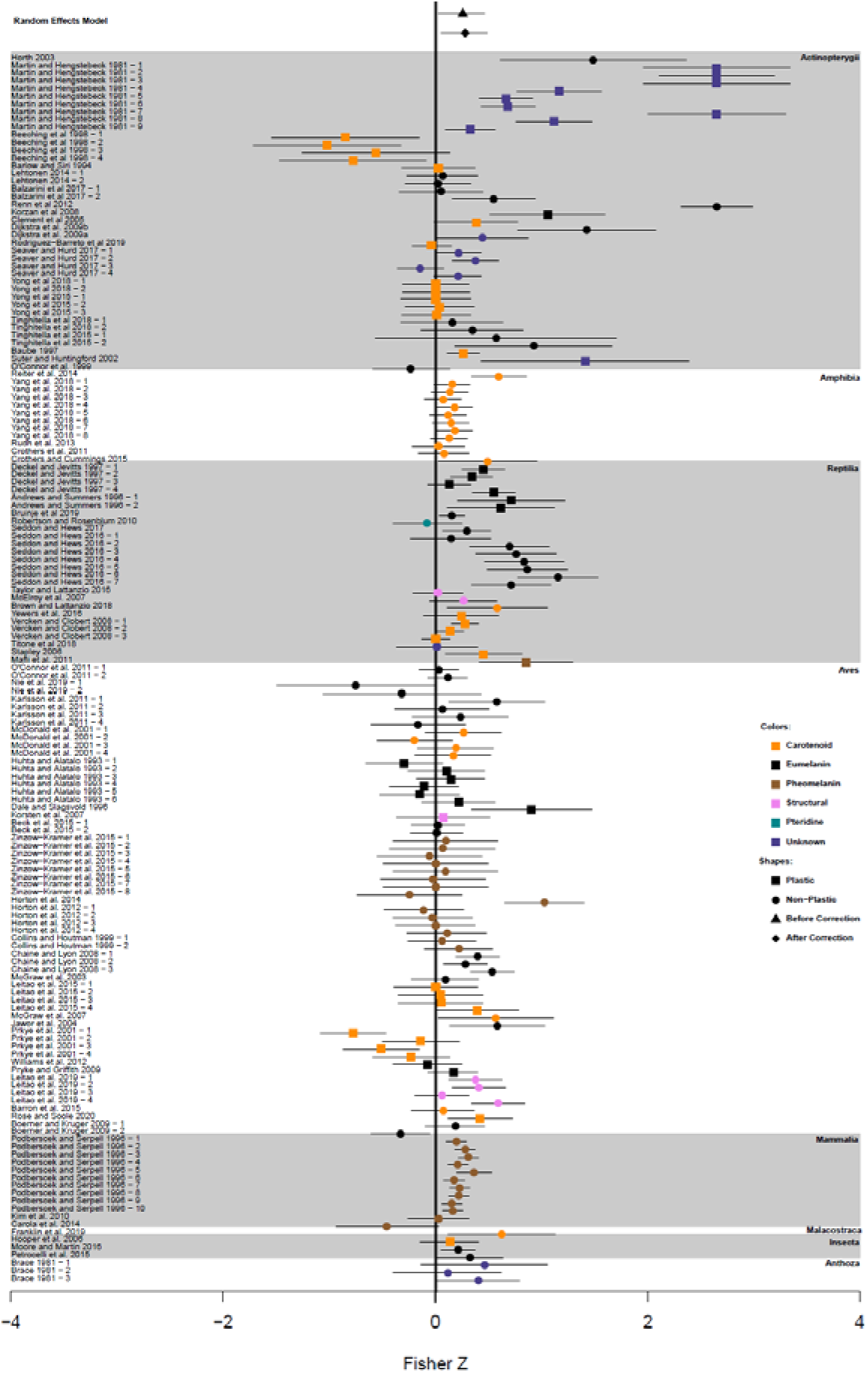
The effect size and 95% credible intervals for all studies listed by family. The study names are on the left side of the graph and any study with a dash then a number is a second effect size from the same study. The colors of the points indicate the color class, while the shape of the point indicates if the trait was plastic or non-plastic. The points at the top of the graph are the mean effect size and 95% credible intervals for the random effects model. The black triangle and line through it indicate the mode mean with 95% credible intervals before publication bias correction. Black diamond and line indicate model means with 95% credible intervals after publication bias correction.

In the model that included all of the random effects, phylogenetic signal accounted for 21.0% of the variation in the dataset (Supplemental Table S2, Figure 2). However, when we removed the phylogeny, species, and study sequentially, the only term that caused substantially poorer fit when removed was the study term (Spiegelhalter et al. 2002; Supplemental Table S2). Removing only the phylogenetic signal or removing the tree alone did not result in a poorer fit when compared to the full random effects model. This suggests that the “phylogenetic” signal arises because some taxonomic groups are represented by one or a very few studies (e.g., Amphibia, see Figure 2). We therefore interpret this signal as mainly reflecting variation among studies, not a true phylogenetic pattern.

None of the fixed effect moderators we investigated improved the fit of the full random effects model; that is, adding any moderator increased the DIC value. Some of the mixed effects models exhibited DIC values that were close to that of the full random effects model (Supplemental Table S2 and Supplemental Table S3). For example, a mixed model that included type of aggressive act as a moderator had a DIC value that was very slightly higher than that of the full random effects model (- 262.442, Supplemental Table S3). No other model had a lower DIC value than these two. Critically, no models that included color class (eumelanin, pheomelanin, carotenoid, structural, or unknown colors) as a moderator, either alone or in combination with any other moderator, ever achieved a DIC value below -258. Consequently, there is no evidence in this data set for color-type differences in the association between intensity of color and aggression.

We did find evidence of a publication bias for the full random effects model. In this model, the intercept for the modified Egger’s regression was significantly different from zero (intercept ± SE: 0.777±0.3815, t-value = 2.03, p-value = 0.0433). The trim and fill method added zero imputed values to the original 159 effect size estimates and indicated an adjustment of 0.0233 to the Fisher Z values to restore funnel plot symmetry (Figure 3). After this adjustment, the overall mean Fisher Z value increased, but still indicated a positive association between color and aggressive behavior (adjusted mean = 0.274 and 95% credible interval = (0.041, 0.481); Pearson’s correlation coefficient adjusted mean = 0.267, and 95% credible interval = (0.041, 0.447); Figure 2).

**Figure 3:**
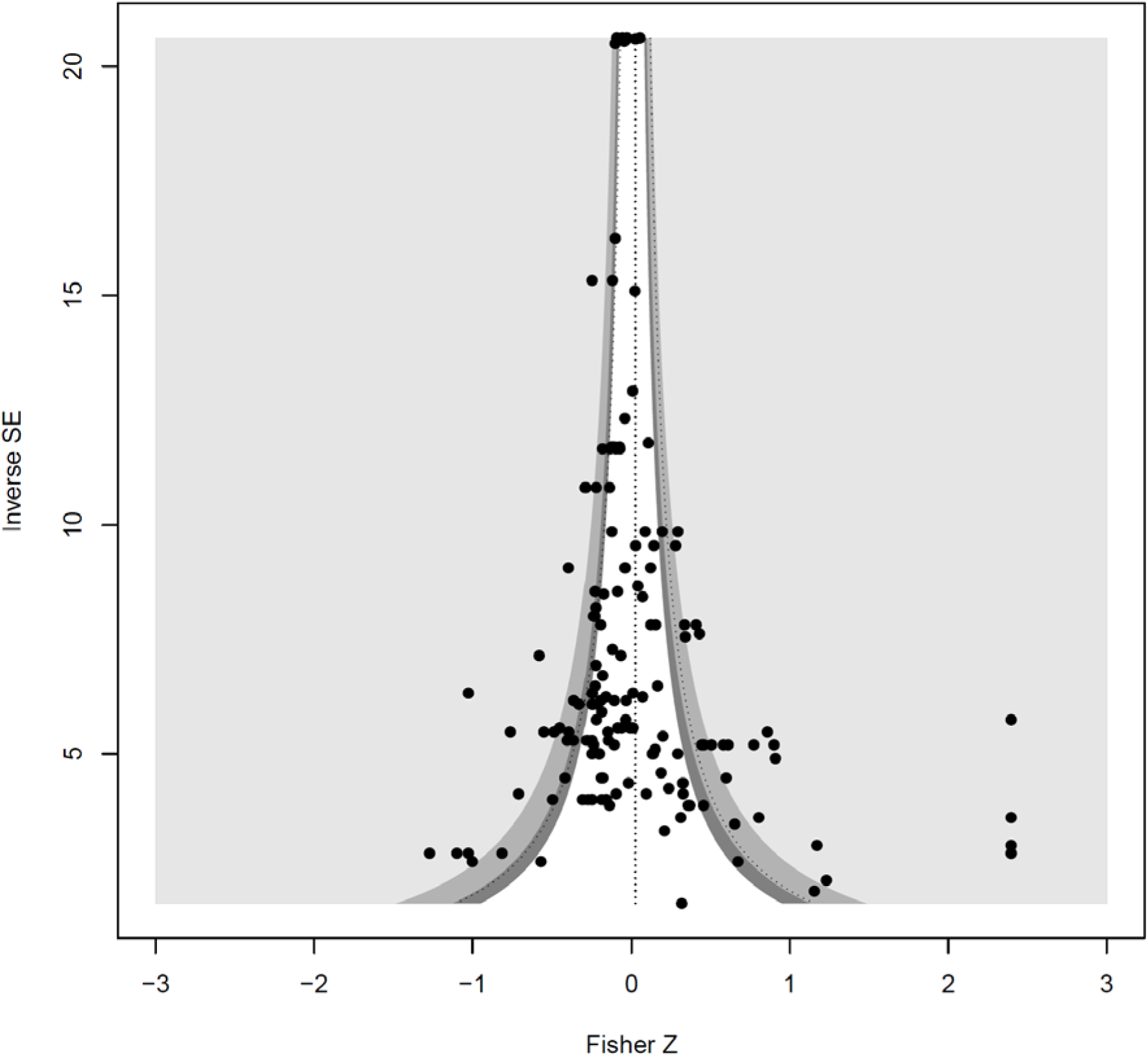
Funnel Plot of Random Effects Only Model. No points were added to restore plot symmetry.

To determine if the melanocortin pleiotropy hypothesis would be better supported if we reclassified “unknown” color types as “eumelanic”, we ran the mixed effects model that included color class as a moderator on this recoded data set. This “recoded” mixed model had a similar posterior effect distribution (mean = 0.261, 95% credible interval: (-0.020, 0.559)). As in the original data set, adding the color moderator did not improve the fit (DIC Value for recoded color model: -259.414, Supplemental Table S3). Notably, the mixed model did not provide a better fit than the random effects model for either data set, suggesting that the relationship between color intensity and aggression does not vary based on the type of color, as predicted by the melanocortin pleiotropy hypothesis.

### Analysis of data subsets

Results did not differ substantially in analyses of data subsets that included only those studies that controlled for animal condition, social rank, or age. As in analyses of the full data set, the random effects model was a better fit than a model that included color type as a moderator for all the data subsets (Supplemental Table S4). The mean posterior effect sizes for the relationship between color and aggression was also similar in magnitude to the mean estimate of the full data set, ranging from 0.177 to 0.307 (Supplemental Table S4). Fewer studies controlled for animal age (beyond juvenile/adult) compared to those that controlled for social rank or condition. Analysis of the subset that did control for age (16 studies) or for age, condition, and social rank (9 studies), also produced similar mean posterior effect size estimates, but in these cases, the credible interval for the mean effect size did overlap zero (Supplemental Table S4).

## Discussion

The meta-analysis of 70 published studies indicated a positive association between measures of aggression and measures indicating more colorful individuals. For simplicity, we used the term “colorful” to refer to variation in hue, intensity, or area as used in the original studies. This positive relationship held after correcting for publication bias. No moderator that we evaluated improved the fit of the meta-analytic model to the data, indicating a robust relationship between color and aggression. Critically, we found no evidence that this relationship depended on the type of color (eumelanic, phaeomelanic, carotenoid, or structural), counter to predictions of the melanin-pleiotropy and carotenoid-condition-dependence hypotheses. This result was unchanged when we recoded colors of unknown cause, but for which a melanocortin-based mechanism is plausible. Both the badge-of-status and general condition-dependence hypotheses are consistent with this pattern. However, under general condition-dependence, we expect the effect size to be sensitive to moderators indicating whether and how condition was accounted for in the study design, or to moderators that are plausibly associated with condition, such as social rank and age. None of these moderators explained variation in effect sizes in our analyses, and none affected the (lack of) dependence of effect size on type of color.

It is possible that different mechanisms underly similar correlations between aggression and different kinds of coloration. For example, it is possible that covariation between melanin-based colors and aggression in the relevant studies in our data set is indeed regulated by variation in the melanocortin pathway, and that covariation between aggression and carotenoid color is regulated by condition dependence that was not accounted for by any of the proxies for condition that we included in our analyses. However, the badge of status hypothesis is a more parsimonious explanation for these patterns (Tibbetts and Dale 2004; Santos et al. 2011).

Potential caveats to this conclusion include failure of moderators to improve the fit can result from including studies with low sample size and high variation, or from including few studies assessing the moderators of interest (Ginzburg and Jensen 2004; Lajeunesse 2009). For key moderators, this study had robust sample sizes. For example, in the condition category, 101 measures derived from studies that accounted for animal condition in some way, while 58 measures did not. In the subset of data that included only studies accounting for condition in some way, 34 measures accounted for both length and weight, 20 for weight only, and 47 for length only. In addition, the low heterogeneity observed in the pure random-effects model suggests that power was not severely compromised by low sample size or high uncertainty in effect size estimates. Nevertheless, some combinations of moderators (e.g., color class, social rank, and age) were represented by few effect sizes in our data set (Supplemental Table S4). Consequently, if condition is affected by adult age, as seems likely, current literature might be inadequate to evaluate the general-condition-dependence hypothesis. We therefore suggest that future studies control for animal age, body condition, and social rank (where applicable).

We did observe considerable variation in effect sizes among studies, and the among-study variation was the main contributor to heterogeneity in our analysis. In contrast, species and phylogenetic relatedness did not explain additional heterogeneity. For example, the most extreme effect sizes (both positive and negative) are found within Actinopterygii (Figure 2). The strongest negative association derived from a single study of orange color intensity in female convict cichlids (Beeching et al. 1998). In that study, females with the most orange color displayed the lowest level of aggression toward a stimulus female. The most positive associations between color and aggression occurred in a study of plastic eye color in juvenile guppies (*Poecilia reticulata*; Martin and Hengstebeck 1981). Fish with darker eye color engaged in and won more aggressive encounters than the light-eyed fish. Losers in these encounters also lightened their eye color, consistent with some studies of traits deemed to be badges of status (Dey et al. 2014).

In our data set, effect sizes within studies tended to be tightly clustered. This clustering suggests that even closely-related species can differ appreciably in the associations we measured. Alternately, this pattern could be driven by differences in methodologies, as was found in a meta-analysis of birds (Santos et al. 2011). We categorized studies based on whether measures of aggression were direct or indirect (as in Santos et al. 2011), and on whether the general methodology was observational or experimental; neither of these categorizations was associated with variation in effect size. We do recommend that future studies ideally take an experimental approach and that they include direct measures of aggression when possible.

Of the four hypotheses explaining consistent associations between animal coloration and aggression, our results are most parsimoniously explained by the badge of status hypothesis. This hypothesis proposes that aggressiveness, fighting ability, or dominance status is honestly reflected by variation in a trait that is perceptible to conspecifics (Rohwer 1975; McGraw et al. 2003; Tibbetts and Dale 2004). While many studies suggest that melanocortin-based genetic pleiotropy can constrain the joint evolution of color and behavior, whether the badge-of-status mechanism should impose such constraints has received less attention. Under the badge-of-status hypothesis, the correlation between color and aggression could be caused by pleiotropy (Santos et al. 2011; Küpper et al. 2016; Lamichhaney et al. 2016; Sánchez-Tójar et al. 2018). The supergene regulating feather coloration and social status in ruffs is a prime example (Küpper et al. 2016; Lamichhaney et al. 2016). The relationship between head stripe color and aggression in white-throated sparrows is another likely example of genetically-based covariation that arises from the badge of status mechanism (Lowther 1961; Knapton and Falls 1983; Tuttle 2003; Tuttle et al. 2016; Hedrick et al. 2018). Variation in a badge of status could also be regulated by non-genetic variation in resource availability or acquisition, variation in exposure to disease or parasites, or non-genetic maternal effects (Rohwer 1975; Dawkins and Krebs 1978). However, like for other condition-dependent mechanisms, our analysis does not support this interpretation for the overall trend we observed: a moderate positive correlation between coloration and behavior across *Animalia*. We therefore propose that this moderate correlation is underlain by genetic covariation between behavior and color traits that serve as badges of status. A key question remains: is this genetic correlation underlain by the same mechanism across different color types? This could occur, for example, if genetic variation in condition regulates both behavior and color, irrespective of the color of the badge. Alternatively, do different mechanisms, e.g. melanocortin pleiotropy in the case of eumelanin color, and genetic variation in condition for carotenoid colors. Studies of the basis of genetic covariance variety of systems will be needed to answer this question.

To date, few studies have attempted to partition covariation between color and behavior into heritable and non-heritable components, and most of these have been in systems characterized by discrete color polymorphisms (e.g., Tuttle 2003; Küpper et al. 2016; Lamichhaney et al. 2016). In tractable organisms (mainly short-lived invertebrates), artificial selection or experimental evolution could be used to measure genetic correlation between traits and thus, directly assess whether color-behavior associations impose constraints on evolutionary change. Many free-living populations (mainly vertebrates) have been the subject of long-term observational study, and now have pedigree data that could allow genetic correlations between color and behavior to be estimated (Clutton-Brock and Pemberton 2004; Blondel et al. 2006; Charmantier et al. 2006; Foerster et al. 2007; McAdam et al. 2007). These kinds of data will become increasingly available as more long-term studies incorporate genomic analyses of relatedness. Consequently, both experimental and non-experimental studies in a variety of species will be necessary to understand the extent to which color-behavior correlations influence evolutionary dynamics.

## Acknowledgements

We thank the Hughes Lab, Emily DuVal, and David Houle for their helpful comments throughout the data collection process and for valuable suggestions that improved this manuscript.

## Funding Sources

This work was funded by the National Science Foundation Grant # DEB-1740466.

## Author Contribution

SNR: Conceptualization, Methods development/experimental design, Data Collection, Data Analysis, Data Visualization, Model Analysis, Coding Simulation, Supervision, Writing – original draft and review & editing. EAH: Conceptualization, Methods development/experimental design, Data Collection, Writing – original draft and review & editing. LM, IP, and MP: Data Collection. KAH: Conceptualization, Methods development/experimental design, Supervision, Writing – original draft and review & editing

## Notes

https://github.com/sruckman/meta-analysis

### Competing Interest Statement

The authors have declared no competing interest.

### Summary of Updates

Introduction and Discussion were updated to clarify the badge of status hypothesis and how it relates to our results.

