## Supplemental Figures for "Assessing the role of pleiotropy in the evolution of animal color and behavior: a meta-analysis of experimental studies"

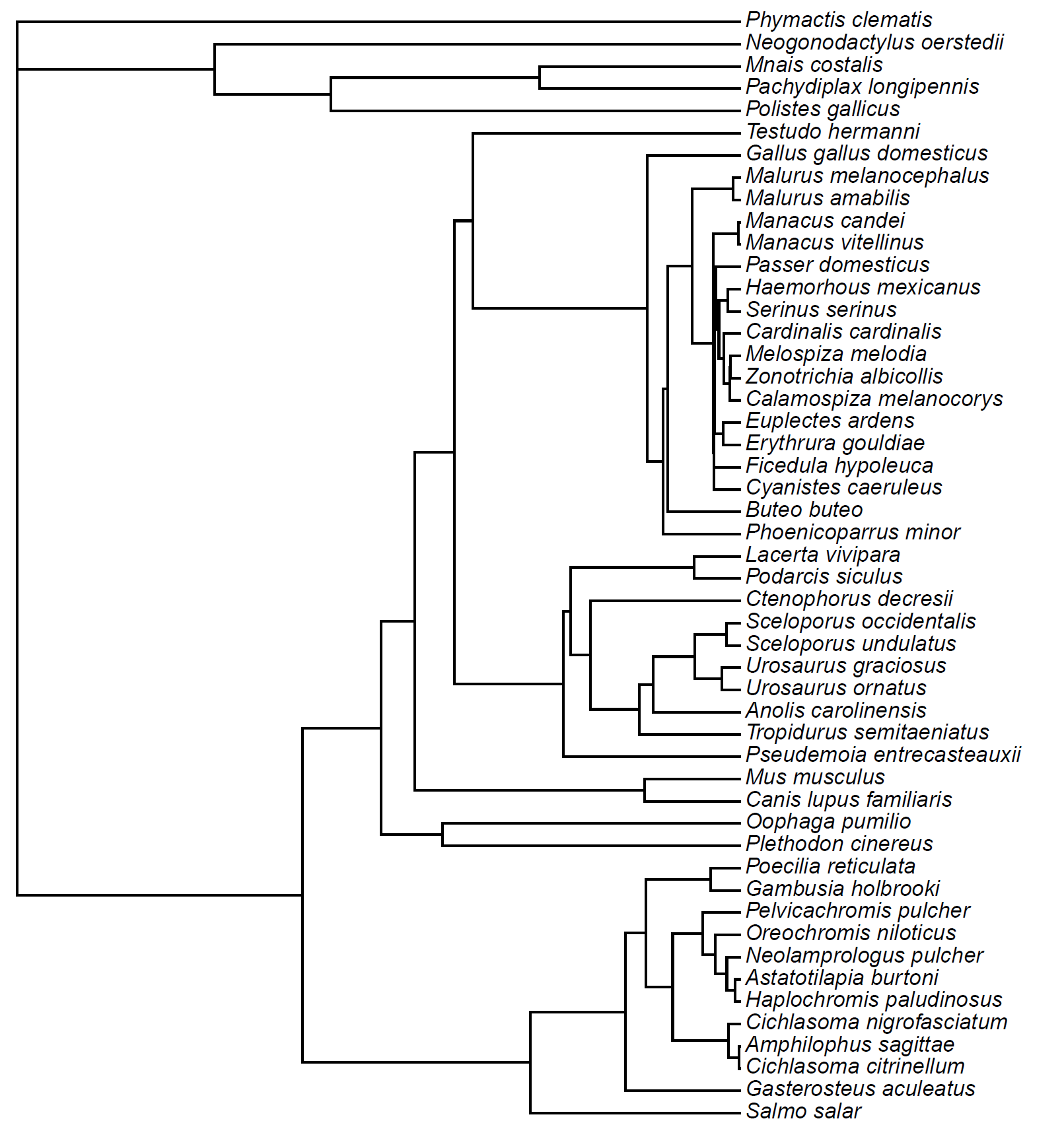

Supplemental Figure S1: Phylogeny for Papers Examined. The phylogeny is an ultrametric tree that was fully resolved to the species level, which we obtained from TimeTree.org.
