## Supplemental Table S1 for "Assessing the role of pleiotropy in the evolution of animal color and behavior: a meta-analysis of experimental studies"

Supplemental Table S1: Body Color Classification Justification. Every species included in the meta-analysis is listed, along with categories of the colors used by the study authors (columns 1-3). Justification for the classification of the colors used in the meta-analysis are in columns 6-7.

| **Species** | **Light Color** | **Dark Color** | **Plastic** | **Breeding** | **Classification** | **Explanation** |
| --- | --- | --- | --- | --- | --- | --- |
| *Amphilophus sagittae* | Gold | Black/dark | No | Yes | Melanin  (eumelanin) | Melanophore death = gold and black is due to melanophore presence (Henning et al., 2010, 2013) |
| *Anolis carolinensis* | Light green | Dark green or Brown | Yes | No | Melanin (eumelanin) | Melanophore location under the xanthophores. Below = green and above = brown (Taylor and Hadley, 1970) |
| *Astatotilapia burtoni* | Blue | Yellow | Yes | No | Melanocortin (eumelanin) | Exogenous α-melanocyte-stimulating hormone (α-MSH) increases yellowness of the body and dispersal of xanthophore pigments in both morphs (Dijkstra et al., 2017) |
| *Astatotilapia burtoni* | No black | Black | No | No | Melanocortin (eumelanin) | Eumelanin = black bars on males (Li et al., 2021) |
| *Buteo Buteo* | Light white | Dark/brown | Yes | No | Melanocortin (eumelanin) | Eumelanic morphs (Chakarov et al., 2008) |
| *Calamospiza melanocorys* | Light black | Dark black | No | Yes | Melanocortin (eumelanin) | The black breeding plumage of males results from melanin deposition during feather growth (Chaine). |
| *Canis lupus familiaris* | White | Fawn | No | No | Melanocortin (pheomelanin) | Fawn is based on the agouti gene and is based on pheomelanin (Schmutz and Berryere, 2007) |
| *Canis lupus familiaris* | Black | Red/Gold | No | No | Melanocortin (pheomelanin) | E gene = pheomelanin when red (Schmutz and Berryere, 2007) |
| *Cardinalis cardinalis* | Light black | Dark black | No | No | Melanocortin (eumelanin) | (Jawor and Breitwisch, 2003) |
| *Cichlacoma citrinellum* | Grey | Golden | Yes | No | Carotenoid | No melanophores in gold only carotenoids (Webber et al., 1973) |
| *Cichlasoma nigrofasciatum* | Dull orange | Dark orange | Yes | No | Carotenoid | Carotenoids produce brighter orange (Barlow, 1976) |
| *Ctenophorus decresii* | Grey/Yellow | Orange | Yes | No | Carotenoid | Yellow is generated primarily by carotenoids and orange is generated by the combination of carotenoids and drosopterin (pteridine) (Rankin et al., 2016) |
| *Cyanistes caeruleus* | Less blue | Bluer | Yes | No | Structural | Crown of the blue tit is structural colors and changes over seasons (Delhey et al., 2010) |
| *Erythrura gouldiae* | Red | Black | Yes | No | Melanocortin (eumelanin) | Recessive alleles produce eumelanin (black melanin), which masks the effects of the carotenoids responsible for red/yellow headedness and produces the black-headed morph (Brush and Seifried, 1968) |
| *Euplectes ardens* | Yellow/Orange | Red | Yes | No | Carotenoid | (Prager and Andersson, 2010) |
| *Ficedula hypoleuca* | Dull brown | Dark brown | Yes | No | Melanocortin (eumelanin) | More eumelanin pigment (responsible for darker colors) than pheomelanin (which is involved in browns) (Potti et al., 2014). |
| *Gallus gallus domesticus* | White | Wild type (red) | No | No | Melanocortin (eumelanin) | Dark red pigment = eumelanin (Nätt et al., 2007) |
| *Gambusia holbrooki* | Silver | Black | No | No | Melanocortin (eumelanin) | (Angus, 1989) or temperature dependent (Horth, 2003) |
| *Gasterosteus aculeatus* | Dull or no red | Bright red | Yes | Yes | Carotenoid | (Brush and Reisman, 1965) |
| *Gasterosteus aculeatus* | Red | Black | No | No | Melanocortin (eumelanin) | (Lewandowski and Boughman, 2008) |
| *Haemorhous mexicanus* | Light red | Dark red | No | No | Carotenoid | (Lendvai et al., 2013) |
| *Haplochromis paludinosus* | Red | Blue | No | No | Unknown | Not described |
| *Haplochromis paludinosus* | Plain (orange) | Black | No | No | Melanocortin (eumelanin) | (Dijkstra et al., 2009) |
| *Lacerta vivipara* | Orange | Yellow | Yes | No | Carotenoid | Carotenoids reflect genetically based color morphs, but iridophores lead to plasticity in color (San-Jose et al., 2013) |
| *Luscinia cyanura* | Brown | Blue | No | No | Structural | (Morimoto et al., 2006) |
| *Malurus amabilis* | Grey/less blue | Blue | No | Yes | Structural | (Fan et al., 2019) |
| *Malurus melanocephalus* | Brown | Red/Black | No | Yes | Carotenoid | (Khalil et al., 2020) |
| *Manacus candei* | White | Golden | No | Yes | Carotenoid | (Bennett et al., 2021) |
| *Manacus vitellinus* | White/lemon | Golden | No | Yes | Carotenoid | (Bennett et al., 2021) |
| *Melospiza melodia* | Light brown | Dark brown | No | Yes | Melanocortin (eumelanin) | Darker = eumelanin and brown = pheomelanin (Beck et al., 2018) |
| *Mnais costalis* | Clear | Orange | Yes | No | Carotenoid | (Maoka et al., 2020) |
| *Mus musculus* | Non-agouti | Agouti | No | No | Melanocortin (pheomelanin) | (Lamoreux et al., 2001) |
| *Neogonodactylus oerstedii* | Light meral spots | Dark meral spots | No | No | Carotenoid | Carotenoproteins create the spots (Franklin et al., 2019) |
| *Neolamprologus pulcher* | No black stripes | Black stripes | No | No | Melanocortin (eumelanin) | (Balzarini et al., 2017) |
| *Oophaga pumilio* | Light red | Dark red | No | No | Carotenoid | (Rodríguez et al., 2020) |
| *Oophaga pumilio* | Green | Red | No | No | Carotenoid | (Rodríguez et al., 2020) |
| *Oophaga pumilio* | Red | Blue | No | No | Carotenoid | Carotenoid leads to blue skin (Rodríguez et al., 2020) |
| *Oreochromis niloticus* | Light (yellow) | Darker (red) | Yes | No | Carotenoid | Diet carotenoids lead to color differences (Wang et al., 2021) |
| *Pachydiplax longipennis* | Less black | Blacker | No | No | Melanocortin (eumelanin) | Most likely eumelanin (Moore and Martin, 2016) |
| *Passer domesticus* | Less black | Blacker | No | No | Melanocortin (eumelanin) | Black eumelanin based feathers (Rojas Mora et al., 2016) |
| *Pelvicachromis pulcher* | Yellow | Red | No | No | Unknown | Not described. May be carotenoids since it’s a fish. |
| *Phoenicoparrus minor* | White | Pink | Yes | No | Carotenoid | Dietary carotenoids (Fox et al., 1967) |
| *Phymactis clematis* | Green | Red | No | No | Unknown | Not described. |
| *Plethodon cinereus* | Unstriped black | Striped red | No | No | Carotenoid | Erythrophores present in red backed but not black backed forms (Bagnara and Taylor, 1970) |
| *Podarcis siculus* | White | Green | No | No | Unknown | Not described. |
| *Poecilia reticulata* | Less black eye | Darker eyes | Yes | No | Unknown | Not described. |
| *Polistes gallicus* | Less black | Blacker | No | No | Melanocortin (eumelanin) | Clypeus spots (most likely eumelanin) (Tibbetts and Dale, 2004) |
| *Pseudemoia entrecasteauxii* | White | Orange | Yes | No | Carotenoid | Most likely carotenoids (López et al., 2009) |
| *Salmo salar* | Pale/grey | Dark grey | No | No | Melanocortin (eumelanin) | Darker spots due to eumelanin (Kittilsen et al., 2009) |
| *Salmo salar* | Light eyes | Dark eyes | Yes | No | Unknown | Not described. |
| *Sceloporus occidentalis* | White | Black | No | No | Melanocortin (eumelanin) | (Seddon and Hews, 2016) |
| *Sceloporus undulatus* | Light orange | Dark orange | No | No | Pteridine | Not carotenoids but instead pteridines lead to orange coloration (Morrison et al., 1995) |
| *Serinus serinus* | Less yellow | More yellow | Yes | No | Carotenoid | Carotenoids lead to yellow coloration (Stradi et al., 1995) |
| *Testudo hermanni* | Light brown | Dark brown | Yes | No | Melanocortin (pheomelanin) | Turtles with brown shells produce pheomelanin (Roulin et al., 2013) |
| *Tropidurus semitaeniatus* | Yellow | Black | No | Yes | Melanocortin (eumelanin) | Do not say if it is eumelanin or not but is most likely eumelanin because of other lizard species (Bruinjé et al., 2019) |
| *Urosaurus graciosus* | Yellow | Orange | No | No | Carotenoid | Related species orange morph due to carotenoids (Haisten et al., 2015) |
| *Urosaurus ornatus* | Yellow/orange | Blue | No | No | Structural | Related species blue morph due to iridophores (Haisten et al., 2015) |
| *Zonotrichia albicollis* | White | Tan | No | No | Melanocortin (pheomelanin) | Reddish brown feathers = pheomelanin (Morrow and Morrow, 2020) |
