## Supplemental Table S2 for "Assessing the role of pleiotropy in the evolution of animal color and behavior: a meta-analysis of experimental studies"

Supplemental Table S2: Table of Model DIC Values for Random Effects Models.

| **Model** | **Fixed Effects** | **DIC** | **Percent Heterogeneity** | | | **Phylogenetic Signal (H^2^)** |
| --- | --- | --- | --- | --- | --- | --- |
|  |  |  | **Study**  $\boldsymbol{(I}_{\boldsymbol{s}}^{\boldsymbol{2}}\boldsymbol{*100)}$ | **Species**  $\boldsymbol{(I}_{\boldsymbol{u}}^{\boldsymbol{2}}\boldsymbol{*100)}$ | **Total (Percentage)** |  |
| Random Includes: species, study, weights, and tree | None (Significant intercept value) | -263.307 | 2.252 | 0.642 | 2.894 | 0.210 |
| Random  Includes: species, study, and weights Missing: Tree | None | -262.193 | -- | -- | -- | -- |
| Random  Includes: species, tree, and weights Missing: study | None | -245.689 | -- | -- | -- | -- |
| Random  Includes: study and weights  Missing: species and tree | None | -262.540 | -- | -- | -- | -- |
| Random  Includes: species, tree, and study Missing: weights | None | 171.219 | -- | -- | -- | -- |
