## Supplemental Table S3 for "Assessing the role of pleiotropy in the evolution of animal color and behavior: a meta-analysis of experimental studies"

Supplemental Table S3: Table of Model DIC Values for Mixed Effects Models.

| **Model** | **Fixed Effects** | **DIC** | **Percent Heterogeneity** | | | **Phylogenetic Signal (H^2^)** | **Mean and 95% Credible Interval** |
| --- | --- | --- | --- | --- | --- | --- | --- |
|  |  |  | **Study**  $\boldsymbol{(I}_{\boldsymbol{s}}^{\boldsymbol{2}}\boldsymbol{*100)}$ | **Species**  $\boldsymbol{(I}_{\boldsymbol{u}}^{\boldsymbol{2}}\boldsymbol{*100)}$ | **Total (%)** |  |  |
| Mixed | Color Class (full data) | -258.406 | 2.086 | 0.893 | 2.979 | 0.284 | Mean: 0.259  (-0.032, 0.583) |
| Mixed | Color Class (unknown categorized as eumelanin) | -259.414 | 2.102 | 0.925 | 3.027 | 0.290 | Mean: 0.258  (-0.755, 1.277) |
| Mixed | Aggression Measure | -262.442 | 2.331 | 0.525 | 2.856 | 0.174 | Mean: 0.252  (0.059, 0.481)  Direct: 0.309, (0.125, 0.542)  Indirect: 0.18,  (-0.118, 0.416) |
| Mixed | Age Controlled | -256.569 | -- | -- | -- | -- | -- |
| Mixed | Plasticity | -261.145 | -- | -- | -- | -- | -- |
| Mixed | Sex | -260.035 | -- | -- | -- | -- | -- |
| Mixed | Vert/Invert | -260.797 | -- | -- | -- | -- | -- |
| Mixed | Location | -260.208 | -- | -- | -- | -- | -- |
| Mixed | Season | -253.636 | -- | -- | -- | -- | -- |
| Mixed | Life Stage | -259.969 | -- | -- | -- | -- | -- |
| Mixed | Geography | -261.216 | -- | -- | -- | -- | -- |
| Mixed | Social Rank | -257.187 | -- | -- | -- | -- | -- |
| Mixed | Obs vs Exp | -261.297 | -- | -- | -- | -- | -- |
| Mixed | Condition | -259.877 | -- | -- | -- | -- | -- |
| Mixed | Plasticity and Vert/Invert | -260.332 | -- | -- | -- | -- | -- |
| Mixed | Color Class and Life Stage | -257.861 | -- | -- | -- | -- | -- |
| Mixed | Plasticity and Life Stage | -259.957 | -- | -- | -- | -- | -- |
| Mixed | Sex and Life Stage | -259.063 | -- | -- | -- | -- | -- |
| Mixed | Vert/Invert and Life Stage | -259.296 | -- | -- | -- | -- | -- |
| Mixed | Location and Life Stage | -259.654 | -- | -- | -- | -- | -- |
| Mixed | Season and Life Stage | -254.346 | -- | -- | -- | -- | -- |
| Mixed | Plasticity and Sex | -259.487 | -- | -- | -- | -- | -- |
| Mixed | Color Class and Vert/Invert | -258.429 | -- | -- | -- | -- | -- |
| Mixed | Color Class and Location | -257.900 | -- | -- | -- | -- | -- |
| Mixed | Color Class and Season | -251.133 | -- | -- | -- | -- | -- |
| Mixed | Color Class and Plasticity | -257.797 | -- | -- | -- | -- | -- |
| Mixed | Color Class and Sex | -257.166 | -- | -- | -- | -- | -- |
| Mixed | Plasticity and Location | -261.039 | -- | -- | -- | -- | -- |
| Mixed | Plasticity and Season | -252.112 | -- | -- | -- | -- | -- |
| Mixed | Vert/Invert and Sex | -259.306 | -- | -- | -- | -- | -- |
| Mixed | Sex and Location | -259.047 | -- | -- | -- | -- | -- |
| Mixed | Sex and Season | -252.897 | -- | -- | -- | -- | -- |
| Mixed | Vert/Invert and Location | -259.423 | -- | -- | -- | -- | -- |
| Mixed | Vert/Invert and Season | -253.587 | -- | -- | -- | -- | -- |
| Mixed | Location and Season | -254.240 | -- | -- | -- | -- | -- |
| Mixed | Color Class and Aggression Measure | -259.663 | -- | -- | -- | -- | -- |
| Mixed | Color Class and Geography | -257.900 | -- | -- | -- | -- | -- |
| Mixed | Color Class and Social Rank | -256.130 | -- | -- | -- | -- | -- |
| Mixed | Color Class and Obs. vs. Exp. | -258.149 | -- | -- | -- | -- | -- |
| Mixed | Color Class and Condition | -258.124 | -- | -- | -- | -- | -- |
| Mixed | Aggression Measured and Plasticity | -262.165 | -- | -- | -- | -- | -- |
| Mixed | Aggression Measure and Sex | -260.907 | -- | -- | -- | -- | -- |
| Mixed | Aggression Measure and Vert/Invert | -261.304 | -- | -- | -- | -- | -- |
| Mixed | Aggression Measure and Location | -261.519 | -- | -- | -- | -- | -- |
| Mixed | Aggression Measure and Season | -254.748 | -- | -- | -- | -- | -- |
| Mixed | Aggression Measure and Age | -261.151 | -- | -- | -- | -- | -- |
| Mixed | Aggression Measure and Geography | -262.629 | -- | -- | -- | -- | -- |
| Mixed | Aggression Measure and Social Rank | -259.384 | -- | -- | -- | -- | -- |
| Mixed | Aggression Measure and Obs. vs. Exp | -262.061 | -- | -- | -- | -- | -- |
| Mixed | Aggression Measure and Condition | -259.831 | -- | -- | -- | -- | -- |
| Mixed | Plasticity and Geography | -261.075 | -- | -- | -- | -- | -- |
| Mixed | Plasticity and Social Rank | -257.651 | -- | -- | -- | -- | -- |
| Mixed | Plasticity and Obs. vs. Exp | -261.129 | -- | -- | -- | -- | -- |
| Mixed | Plasticity and Condition | -260.062 | -- | -- | -- | -- | -- |
| Mixed | Sex and Geography | -260.167 | -- | -- | -- | -- | -- |
| Mixed | Sex and Social Rank | -257.662 | -- | -- | -- | -- | -- |
| Mixed | Sex and Obs. vs. Exp | -259.393 | -- | -- | -- | -- | -- |
| Mixed | Sex and Condition | -258.770 | -- | -- | -- | -- | -- |
| Mixed | Vert/Invert and Geography | -260.714 | -- | -- | -- | -- | -- |
| Mixed | Vert/Invert and Social Rank | -256.714 | -- | -- | -- | -- | -- |
| Mixed | Vert/Invert and Obs. vs. Exp | -260.496 | -- | -- | -- | -- | -- |
| Mixed | Vert/Invert and Condition | -259.485 | -- | -- | -- | -- | -- |
| Mixed | Location and Geography | -260.198 | -- | -- | -- | -- | -- |
| Mixed | Location and Social Rank | -256.611 | -- | -- | -- | -- | -- |
| Mixed | Location and Obs. vs. Exp | -261.154 | -- | -- | -- | -- | -- |
| Mixed | Location and Condition | -258.860 | -- | -- | -- | -- | -- |
| Mixed | Season and Geography | -253.941 | -- | -- | -- | -- | -- |
| Mixed | Season and Social Rank | -251.773 | -- | -- | -- | -- | -- |
| Mixed | Season and Obs. vs. Exp | -253.971 | -- | -- | -- | -- | -- |
| Mixed | Season and Condition | -253.282 | -- | -- | -- | -- | -- |
| Mixed | Life Stage and Geography | -259.810 | -- | -- | -- | -- | -- |
| Mixed | Life Stage and Social Rank | -256.917 | -- | -- | -- | -- | -- |
| Mixed | Life Stage and Obs. vs. Exp | -260.170 | -- | -- | -- | -- | -- |
| Mixed | Life Stage and Condition | -259.290 | -- | -- | -- | -- | -- |
| Mixed | Geography and Social Rank | -258.682 | -- | -- | -- | -- | -- |
| Mixed | Geography and Obs. vs. Exp | -258.478 | -- | -- | -- | -- | -- |
| Mixed | Geography and Condition | -260.563 | -- | -- | -- | -- | -- |
| Mixed | Social Rank and Obs. vs. Exp | -257.775 | -- | -- | -- | -- | -- |
| Mixed | Social Rank and Condition | -256.129 | -- | -- | -- | -- | -- |
| Mixed | Obs. vs. Exp and Condition | -259.735 | -- | -- | -- | -- | -- |
| Mixed | Age Controlled and Color Class | -252.745 | -- | -- | -- | -- | -- |
| Mixed | Age Controlled and Agg Measure | -256.907 | -- | -- | -- | -- | -- |
| Mixed | Age Controlled and Plasticity | -255.395 | -- | -- | -- | -- | -- |
| Mixed | Age Controlled and Sex | -254.937 | -- | -- | -- | -- | -- |
| Mixed | Age Controlled and Vert/Invert | -256.354 | -- | -- | -- | -- | -- |
| Mixed | Age Controlled and Location | -255.721 | -- | -- | -- | -- | -- |
| Mixed | Age Controlled and Seasonality | -248.600 | -- | -- | -- | -- | -- |
| Mixed | Age Controlled and Life Stage | -255.462 | -- | -- | -- | -- | -- |
| Mixed | Age Controlled and Geographic | -255.101 | -- | -- | -- | -- | -- |
| Mixed | Age Controlled and Social Rank | -252.442 | -- | -- | -- | -- | -- |
| Mixed | Age Controlled and Obs. vs. Exp. | -255.794 | -- | -- | -- | -- | -- |
| Mixed | Age Controlled and Condition | -256.143 | -- | -- | -- | -- | -- |
| Mixed | Color, Plasticity, and Color by Plasticity | -245.081 | -- | -- | -- | -- | -- |
| Mixed | Color, Sex, and Color by Sex | -246.482 | -- | -- | -- | -- | -- |
| Mixed | Sex, Plasticity, and Sex by Plasticity | -253.227 | -- | -- | -- | -- | -- |
