## Supplemental Table S4 for "Assessing the role of pleiotropy in the evolution of animal color and behavior: a meta-analysis of experimental studies"

Supplemental Table S4: Table of Model DIC Values for Subset Models. Random-effects-only models of different data subsets are listed first in the Table, followed by corresponding models that included the color moderator.

| **Model** | **Fixed Effects** | **DIC** | **Number of Papers** | **Number of Effect Sizes** | **Effect Sizes per Color Class** | **Mean and 95% Credible Interval** |
| --- | --- | --- | --- | --- | --- | --- |
| Random Includes: species, study, weights, and tree | None  Subset by: Controlled Social Rank | -212.052 | 53 | 124 | 40 carotenoid, 40 eumelanin, 28 pheomelanin, 1 pteridine, 6 structural, and 9 unknown | Mean: 0.203  (0.048, 0.383) |
| Random Includes: species, study, weights, and tree | None  Subset by: Controlled Condition | -113.775 | 53 | 101 | 39 carotenoid, 48 eumelanin, 1 pheomelanin, 1 pteridine, 6 structural, and 6 unknown | Mean: 0.307  (0.032, 0.599) |
| Random  Includes: species, study, weights, and tree | None  Subset by: Controlled Age | 13.235 | 16 | 25 | 8 carotenoid, 13 eumelanin, 2 pheomelanin, 1 unknown, and 1 structural | Mean: 0.260  (-0.430, 1.078) |
| Random Includes: species, study, weights, and tree | None  Subset by: Controlled Social Rank and Condition | -105.407 | 40 | 78 | 31 carotenoid, 34 eumelanin, 1 pheomelanin, 1 pteridine, 6 structural, and 5 unknown | Mean: 0.233  (0.073, 0.388) |
| Random  Includes: species, study, weights, and tree | None  Subset by: Controlled Age, Social Rank, and Condition | -4.436 | 9 | 15 | 8 carotenoid, 1 unknown, 5 eumelanin, and 1 structural | Mean: 0.177  (-0.181, 0.523) |
| Mixed | Color Class  Subset by: Controlled Social Rank | -207.236 | 53 | 124 | 40 carotenoid, 40 eumelanin, 28 pheomelanin, 1 pteridine, 6 structural, and 9 unknown | Mean: 0.216,  (-0.046, 0.486) |
| Mixed | Color Class  Subset by: Controlled Condition | -111.167 | 53 | 101 | 39 carotenoid, 48 eumelanin, 1 pheomelanin, 1 pteridine, 6 structural, and 6 unknown | Mean: 0.314  (-0.078, 0.730) |
| Mixed | Color Class Subset by: Controlled Age | 12.208 | 16 | 25 | 8 carotenoid, 13 eumelanin, 2 pheomelanin, 1 unknown, and 1 structural | Mean: 0.289  (-1.225, 1.844) |
| Mixed | Color Class  Subset by: Controlled Social Rank and Condition | -102.804 | 40 | 78 | 31 carotenoid, 34 eumelanin, 1 pheomelanin, 1 pteridine, 6 structural, and 5 unknown | Mean: 0.239  (0.007, 0.482) |
